# uORF4u: a tool for annotation of conserved upstream open reading frames

**DOI:** 10.1101/2022.10.27.514069

**Authors:** Artyom A. Egorov, Gemma C. Atkinson

## Abstract

**Summary:** Upstream open reading frames (uORFs, encoding so-called leader peptides) can regulate translation and transcription of downstream main ORFs (mORFs) in prokaryotes and eukaryotes. However, annotation of novel functional uORFs is challenging due their short size of usually less than 100 codons. While transcription- and translation-level next generation sequencing (NGS) methods can be used for genome-wide uORF identification, this data is not available for the vast majority of species with sequenced genomes. At the same time, the exponentially increasing amount of genome assemblies gives us the opportunity to take advantage of evolutionary conservation in our predictions of ORFs.

Here we present a tool for conserved uORF annotation in 5′ upstream sequences of a user-defined protein of interest or a set of protein homologues. It can also be used to find small ORFs within a set of nucleotide sequences. The output includes publication-quality figures with multiple sequence alignments, sequence logos and locus annotation of the predicted uORFs in graphical vector format.

**Availability and Implementation:** uORF4u is written in Python3 and runs on Linux and MacOS. The command-line interface covers most practical use cases, while the provided Python API allows usage within a Python program and additional customisation. Source code is available from the GitHub page: https://github.com/art-egorov/uorf4u. Detailed documentation that includes an example-driven guide available at the software home page: https://art-egorov.github.io/uorf4u.

## Background

Upstream open reading frames (uORFs, encoding so-called leader peptides) regulate expression downstream genes via translational and/or transcriptional attenuation in bacteria, archaea, eukaryotes and viruses (for review see (1,2)). Furthermore, some small uORFs have been shown to be translated into functional proteins (3,4). Annotation of uORFs is essential for understanding complex regulation mechanisms, including inducible expression of antibiotic resistance genes upon an antibiotic challenge (2,5). However, prediction of uORF is complicated by their properties, such as short length, variable distance to the main ORF (mORF) and unusual sequence composition. A breakthrough in genome-wide annotation of translated regions including uORFs came with the single-nucleotide resolution sequencing method ribosome profiling (Ribo-Seq) (6,7). Several bioinformatics tools were created for annotation of uORFs based on Ribo-Seq data, for example uORF-Tools (8) and uORF-seqr (9). However, Ribo-Seq data is available only for a limited number of usually model organisms, and annotation of short uORFs is often complicated by noise in phased signal tracks that appears due to the stochastic nature and specific cutting preferences of RNases (10). Another limitation of the Ribo-Seq approach to annotation is that translation of uORFs may only be induced in certain environmental or cell conditions; for example, translation of eukaryotic GCN4 uORFs depends on amino acid abundance (11).

In the absence of ribosome profiling data, researchers often annotate uORFs in a manual or semi-manual way. This involves retrieval of the 5’ upstream regions of protein coding genes of interest from sequence databases, and manual inspection of the region, with or without the aid of a sequence alignment to indicate functional conservation. Tell-tale signatures of potential uORFs are the presence of Shine-Dalgarno elements (in the case of prokaryotes), along with start and stop codons in the right context and frame (12-14). However, sequence gazing is a time-consuming and tedious task. Thus, various methods have been developed to automate ORF annotation, even in the absence of expression data. The tool sORF finder (15) takes advantage of nucleotide composition bias and can predict small eukaryotic ORFs within 10-100 amino acids length range. MiPepidis (16), and csORF-finder (17) are a ML-based approaches trained on a limited set of eukaryotic organisms. We have found only one tool, uPEPperoni (18)., that when it was available, implemented conservation analysis for prediction. Importantly, these tools are limited to use with eukaryotic genomes.

Thus, there is currently a lack of a simple tool for uORF prediction in both prokaryotes and eukaryotes that leverages sequence conservation. To fill this gap, we set out to build a tool that also includes the following key properties:

1. Ease of installation and implementation with a command-line interface and Python API for higher customisation.
2. Does not have a requirement to build or download large databases. The tool uses the NCBI API to access the RefSeq database (19) and is therefore always up-to date.
3. Supports various input formats: a user-defined protein as the mORF, set of mORF homologues or nucleotide sequences in FASTA format.
4. Thorough documentation with a home page that contains an example-driven guide and detailed API description.
5. Output that contains publication-ready and editable vector graphics.
6. Can be used for sequences across the tree of life (bacteria, archaea, eukaryotes and viruses).

## The uORF4u workflow

The architecture of the uORF4u workflow is defined on user input (**Fig. 1A**). If the input is a single RefSeq protein accession number, uORF4u performs a BlastP search (20) against the online version of the RefSeq protein database (19). The retrieved list of homologues is saved to be used in the subsequent steps. Alternatively, a list of homologues previously curated by the user can be used as input. This is important for allowing the user to decide the breadth and depth of the search; uORFs may differ in their conservation levels across strains and species and therefore it might be necessary to test different input sets. Using the accession list, uORF4u retrieves the corresponding upstream sequences using the NCBI API as implemented in Biopython (21). For eukaryotes, the upstream region is the complete transcript’s 5′ UTR sequence, and for non-eukaryotic microbes, the upstream region is a user-defined (default 500 nucleotides) length from the mORF start codon. The retrieved nucleotide sequences are saved as intermediate output in FASTA format. These sequences, as well as the list of homologues obtained in the previous step, can also be used as optional input for uORF4 in order to skip the previous steps. It is useful to note that when using nucleotide sequences as input, uORF4u can be used as a general conserved ORF search tool, that is, not necessarily upstream of any particular mORF.

**Fig. 1.**
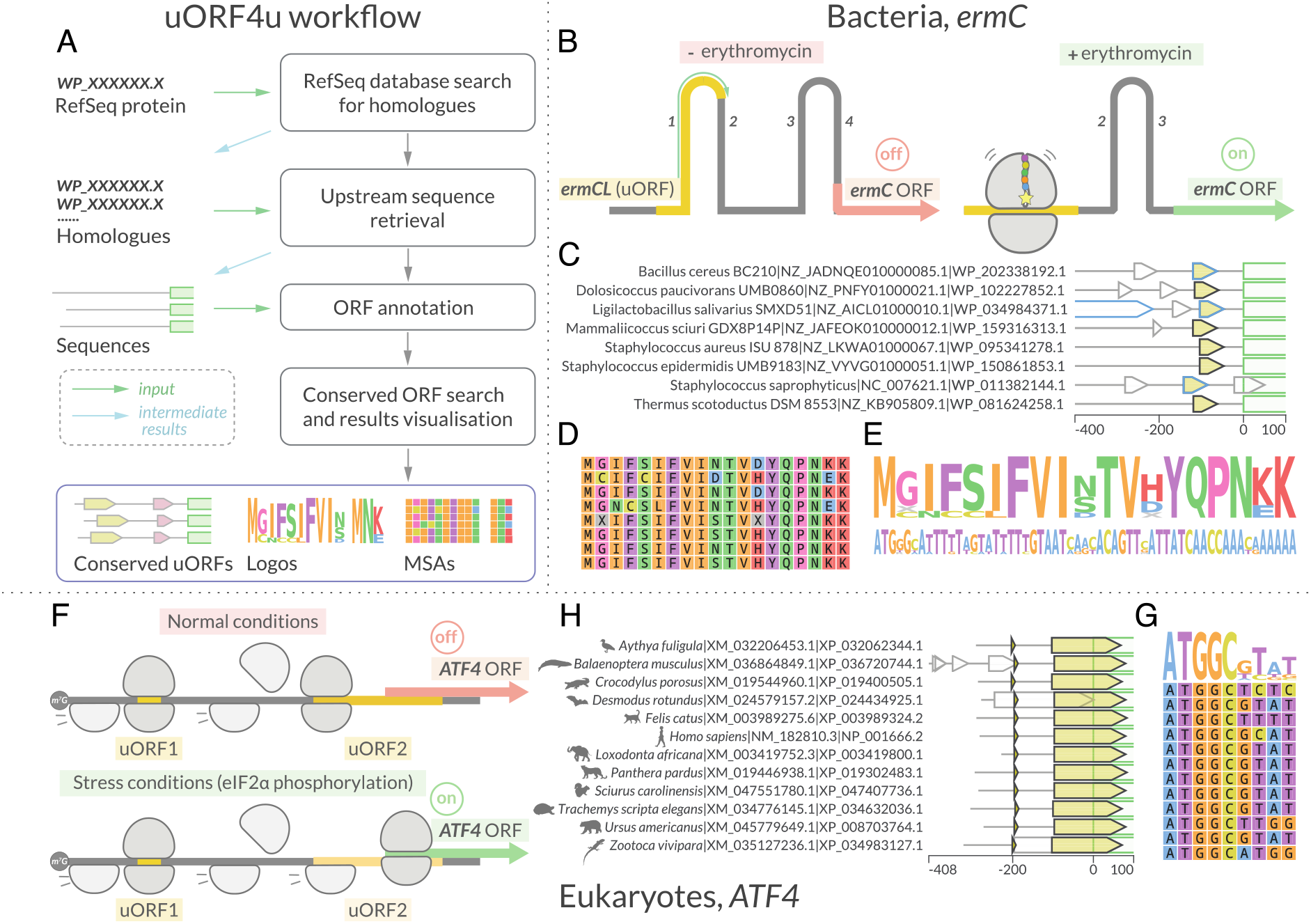
uORF4u workflow and usage examples. (**A**) The uORF4u workflow. The input data is indicated with green arrows. The simplest input is a user-defined RefSeq protein accession number, but could also be a list of homologues (list of RefSeq accession numbers) or a FASTA format file of sequences. Depending on input, uORF4u consists of several steps: search for homologues against the RefSeq database with BlastP (writing the intermediate results (blue arrows) in output), upstream sequence retrieval with results written in an output FASTA file, ORF annotation, and, finally, conservation analysis and results visualisation. (**B**-**E**) Results example using eight bacterial *ermC* proteins as a query (command: *uorf4u -hl WP_202338192*.*1 WP_102227852*.*1 WP_034984371*.*1 WP_159316313*.*1 WP_095341278*.*1 WP_150861853*.*1 WP_011382144*.*1 WP_081624258*.*1 -o ermC -c prokaryotes -annot*. The subset of query proteins was selected from the results obtained with an extended run, see homepage guide for details) (**B**) A bacterial uORF example: regulation of *ermC* expression by translational attenuation. Ribosome stalling on the *ermC* uORF (leader peptide *ermCL*; shown in yellow) is inducible by erythromycin. The arrest alters the regional mRNA structure, exposing the *ermC* SD sequence and allowing translation of the *ermC* mORF (23). (**C**) Annotation plot of the upstream sequences with the conserved *ermCL* uORF. *ermCL* is shown in yellow, the 5′ end of the *ermC* mORF is shown with a green outline, grey outlines indicate other putative ORFs around this locus, and, finally, the blue outline shows ORFs annotated in the RefSeq database. In this case three of the eight were already annotated, and five additional ORFs were annotated by uORF4u (black outline). uORF4u does not use RefSeq ORF annotations in its predictions, and its ability to rediscover both known and missing uORFs validates our strategy (**D, E**) Multiple sequence alignment and sequence logo visualisation of the identified *ermCL* ORFs. (**F-G**) Results example using twelve eukaryotic *ATF4* proteins as a query (command: *uorf4u - hl NP_001666*.*2 XP_036720744*.*1 XP_024434925*.*1 XP_034632036*.*1 XP_008703764*.*1 XP_034983127*.*1 XP_019400505*.*1 XP_003989324*.*2 XP_003419800*.*1 XP_019302483*.*1 XP_047407736*.*1 XP_032062344*.*1 -c eukaryotes*. The same as for ermC example, the list was build based on the extended run results). (**F**) A eukaryotic uORFs example: the expression of *ATF4* (activating transcription factor) is regulated by two uORFs. After translation of the first uORF1, ribosomes are normally able to reinitiate translation at a downstream uORF2 after rebinding the initiating ternary complex *(eIF2-GTP-Met-tR-NA)*. Reduced levels of the ternary complex during stress conditions leads to the ribosome scanning through the uORF2 start codon and instead reinitiating at the *ATF4* uORF (24). (**H**) Annotation plot of the 5’ UTRs with both conserved uORFs shown in yellow with black outline. (**G**) Multiple sequence alignment and sequence logo visualisation of the identified uORF1.

The next step after sequence retrieval is ORF annotation. An ORF is defined as a region between a start codon (alternative start codons can be included as well) and a downstream in-frame stop codon. The minimal length set by default is nine nucleotides (three codons). For prokaryotes this step also includes Shine-Dalgarno (SD) sequence search within a 20-nucleotide window upstream of the start codon. SD sequence annotation is based on the calculation of the SD-antiSD interaction Gibbs free energy (22). For identified potential frames, the tool searches for conserved ORFs using a greedy algorithm: uORF4u iterates through sequences and tries to maximise the sum of pairwise alignment scores between uORFs. The last step in our workflow is generation of multiple sequence alignments (MSA) of the identified conserved uORFs, writing reports and making results visualisation files (annotation plots, sequence logos and MSAs). Examples of output plots are shown on **Fig. 1C-E, H, G**.

## Implementation

uORF4u is written in Python3 and uses multiple python libraries: Biopython (21), configs, argparse, pandas, statistics, Logomaker (25), matplotlib (26), reportlab. Biopython (21) is widely used for different steps; for example, the Blast submodule is used to perform BlastP (20) searching, the Entrez submodule gives access to the NCBI RefSeq database (19), and pairwise alignment between uORFs is performed with Biopython submodules. The Logomaker (25) package API is used to plot sequence logos of conserved ORFs. The MSA is built with MAFFT (v.7505) (27) embedded in the uORF4u library.

The python uORF4u package is available in PyPI (*python3 -m pip install uorf4u*), and the source code is provided on the GitHub page (github.com/art-egorov/uorf4u). Detailed documentation with an installation guide, and an example-driven manual are available at the uORF4u home page (https://art-egorov.github.io/uorf4u/).

The command line interface allows users to run the tool with various standard usage scenarios without any additional effort for user-side scripting. Furthermore, we provide a python API that allows additional customisation.

## Conclusion

The problem of novel uORFs annotation requires specialised tools. Here, we present uORF4u, which performs database parsing, uORF searching, conservation analysis and produces publication-quality images of the results. We believe that as well as identifying uORFs of specific mORFs in targeted analyses our tool paves the way for systematic analysis of uORF genesis, conservation and distribution on the scale of the whole proteomes.

## Acknowledgements

The authors thank Jose Nakamoto and Veda Bojar for uORF4u testing and Vasili Hauryliuk for comments on the manuscript.

